# Decreasing *Wapl* dosage partially corrects transcriptome phenotypes in *Nipbl*-/+ embryonic mouse brain

**DOI:** 10.1101/2022.05.31.493745

**Authors:** Connor M. Kean, Christopher J. Tracy, Apratim Mitra, Matthew T Van Winkle, Claudia M Gebert, Jacob Noeker, Anne L. Calof, Arthur D. Lander, Judith A. Kassis, Karl Pfeifer

**Author notes:** These authors contributed equally to this work (Connor M. Kean, Christopher J. Tracy).

## Abstract

Cohesin rings interact with DNA and modulate expression of thousands of genes. NIPBL loads cohesin onto chromosomes and WAPL takes it off. Heterozygous mutations in *NIPBL* lead to a developmental disorder called Cornelia de Lange syndrome. *Nipbl* heterozygous mice are a good model for this disease but mutations in *WAPL* were not known to cause disease or gene expression changes in mammals. Here we show dysregulation of >1000 genes in *Wapl*^*Δ/+*^ embryonic mouse brains. The patterns of dysregulation are highly similar to *Nipbl* heterozygotes, suggesting that *Wapl* mutations may also cause disease in humans. Since WAPL and NIPBL have opposite effects on cohesin’s association with DNA, we asked whether a heterozygous *Wapl* mutation could correct phenotypes seen in *Nipbl* heterozygous mice. In fact, both gene expression and embryonic growth are partially corrected. Our data are consistent with the view that cohesin dynamics play a key role in regulating gene expression.

## Introduction

The cohesin complex consist of the subunits SMC1, SMC3, RAD21 and Stromalin, which form a ring-like structure that encircles DNA (*1*). Cohesin’s interactions with DNA are dynamic (*2-4*). Cohesin is loaded onto chromosomes by the kollerin complex that consists of Nipped-B-like (NIPBL) and MAU2 (*5*). Once loaded, cohesin can translocate along the chromosome (*4, 6, 7*) or be removed by the releasin complex consisting of PDS5 and WAPL (*8, 9*).

The cohesin complex is required for sister chromatid cohesion and ensures accurate chromosome segregation upon cell division (*10, 11*). Thus, severe disruption of cohesin function results in aneuploidy and cell death. However, studies in Drosophila, zebrafish, mouse, and human reveal that reduced expression of cohesin subunits or of NIPBL alters gene expression and development without evident defects in sister chromatid cohesion and chromosome segregation (*12-16*). For example, Cornelia de Lange syndrome (CdLS) is caused by heterozygous loss-of-function mutations in *NIPBL (17, 18)*. CdLS patients display severe developmental defects that vary significantly from patient to patient but always include neurodevelopmental delays and some degree of intellectual disability (*19, 20*). Mutations in other proteins that alter cohesin function cause similar defects; these developmental defects are collectively known as cohesinopathies (Reviewed in (*1, 21, 22*).

*Nipbl*^*-/+*^ mice effectively phenocopy key features of CdLS (*14, 23*). Late-stage embryos are always smaller than *Nipbl*^*+/+*^ littermates and display an array of developmental defects and organ abnormalities that vary from animal to animal. In a C57BL/6J background (used in this study), heterozygotes die soon after birth. As noted in CdLS patients, cell division in *Nipbl*^*-/+*^ mice appears normal. Instead, mutant phenotypes are associated with changes in gene expression that are typically modest (<2-fold) but occur across hundreds of genes.

Mutations in *NIPBL* account for the majority of CdLS occurrences, and no cases of CdLS have yet been attributed to mutations in *WAPL (24)*. However, compiled data from healthy individuals reveals a dearth of predicted loss of function mutations in *WAPL* coding sequences. Additionally, a single *de novo*, heterozygous, missense mutation in *WAPL* was identified in a patient presenting with neurodevelopmental defects (*25*). Together these findings suggest that *WAPL* heterozygosity, like *NIPBL* heterozygosity, might cause global transcriptional dysregulation.

NIPBL loads cohesin on chromosomes and WAPL removes it. As these two proteins work in opposition, we wondered whether decreasing the dose of *Wapl* could correct phenotypes in *Nipbl*^*-/+*^ mice. Previous studies in Drosophila had shown that decreasing the dosage of *Nipped-B* (the Drosophila homolog of *NIPBL*) could correct a developmental phenotype caused by a dominant-negative *Wapl* allele (*26*). Similarly, reducing *WAPL* function in human cell lines permitted survival of cell lines lacking *NIPBL* and *MAU2 (4)*. Here, we generate and characterize a novel conditional mouse *Wapl* allele. We examined the transcriptomes of *Wapl* heterozygotes and show that like *Nipbl*^*-/+*^ brains, *Wapl*^*Δ/+*^ brains show modest changes in expression levels across hundreds of genes. The genes that are dysregulated in *Wapl*^*Δ/+*^ mostly overlap with genes dysregulated in *Nipbl*^*-/+*^ samples. Our results also show that gene expression changes in *Nipbl*^*-/+*^ mice are typically corrected (at least partially) by decreasing *Wapl* dosage. Similarly, decreasing *Nipbl* dosage partially corrects gene expression changes in *Wapl*^*Δ/+*^ mice. Our results are consistent with a model that cohesin dynamics are important for regulating gene expression (*4*). Finally, we show that decreasing *Wapl* dosage is sufficient to partly rescue *Nipbl*-dependent embryonic growth defects but does not rescue perinatal lethality of *Nipbl*^*-/+*^ pups.

## Results

### *Wapl* loss-of-function is pre-implantation lethal

Wildtype and mutant alleles of *Wapl* are depicted in Fig. 1. Mice carrying a conditional *Wapl* allele (*Wapl*^*Flox*^) were generated as described in Methods. In brief, *loxP* sites were inserted upstream of the *Wapl* promoter and downstream of exon 2. Both homozygous and heterozygous mice carrying this allele are viable and fertile. To generate *Wapl*^*Δ*^ mice, we crossed *Wapl*^*Flox/+*^ males with females homozygous for the *E2a-Cre* transgene (JAX #003724) and then backcrossed progeny to C57BL/6J (JAX #000664). The *Wapl*^*Δ*^ allele was expected to be a null allele based on deletion of both the *Wapl* promoter, the translation initiation site, and the peptide coding sequences in exon 2. *Wapl*^*Δ/+*^ mice were recovered at normal Mendelian frequencies (Table S1). However, as expected both from the essential role of WAPL in chromosome segregation and from previous analyses (*27*), *Wapl*^*Δ/Δ*^ animals were not recovered at weaning (Table S2) or even at the blastocyst stage (Table S3). This shows that *Wapl* function is required for early development in mice.

**Figure 1.**
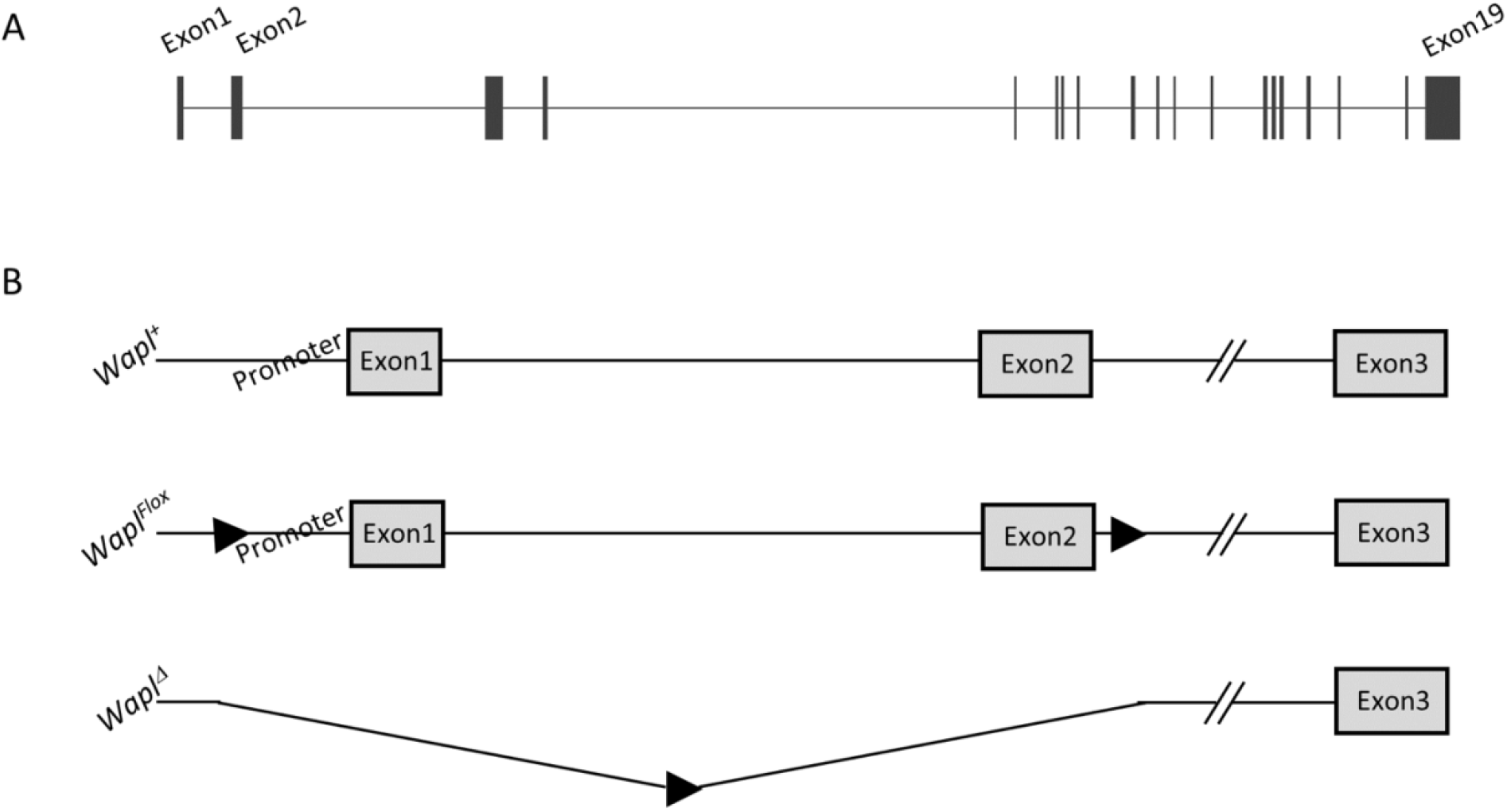
Cartoon depiction of wild type and mutant Wapl alleles. **A**. The *Wapl* gene is encoded on 74,056 bp on mouse chromosome 15. Here we depict the 19 exons of the predominant isoform. For all known isoforms, translation initiates in Exon 2. **B**. Wild type and mutant alleles used in this study. *Wapl*^*Flox*^ carries *loxP* insertions at -517 bp and +3641 bp. (All numbers are relative to the major transcriptional start site). Thus, cre mediated recombination results in deletion of the *Wapl* promoter and exons 1 and 2 to form the *Wapl*^*Δ*^ allele. *LoxP* site, filled arrowhead.

### Decreasing *Wapl* dosage prevents normal mouse development

Although *Wapl*^*Δ/+*^ and *Wapl* ^*Flox/Flox*^ mice are each viable and fertile, we could not generate *Wapl*^*Δ/Flox*^ weanlings. This was tested using two different mating schemes. When *Wapl*^*Δ/+*^ and *Wapl*^*Flox/+*^ mice were intercrossed, no *Wapl*^*Δ/Flox*^ weanlings were identified in 30 progeny (Table S4). Similarly, when *Wapl*^*Δ/+*^ and *Wapl*^*Flox/Flox*^ mice were intercrossed, no *Wapl*^*Δ/Flox*^ weanlings were identified in 20 progeny (Table 1). However, *Wapl*^*Δ/Flox*^ pups are present in expected Mendelian frequencies at embryonic day E17.5 (Table 1). This *Wapl*^*Δ/Flox*^ phenotype is reminiscent of the effect of reducing *Nipbl* gene dosage: *Nipbl*^*-/+*^ heterozygotes are present as late-stage embryos but die before weaning. Altogether, these data show that the *Wapl*^*Flox*^ allele is not completely wildtype and that decreasing *Wapl* gene dosage is detrimental to mouse development.

**Table 1.** *Wapl*^*Δ/Flox*^ weanlings are not viable. χ^2^ = 20.000 with 1 degree of freedom.; P value < 0.001. But *Wapl*^*Δ/Flox*^ E17.5 embryos are found at the expected Mendelian frequency. χ^2^ = 0.077 with 1 degree of freedom; P value = 0.7815.

| Cross: *Wapl*<sup>Δ/+</sup> x *Wapl*<sup>Flox/Flox</sup> |  |  |  |  |  |  |
| --- | --- | --- | --- | --- | --- | --- |
| Collect animals at postnatal day 21 |  |  |  | Collect animals at embryonic day 17.5 |  |  |
| Genotype | *Wapl*<sup>ΔFlox</sup> | *Wapl*<sup>+/Flox</sup> |  |  | *Wapl*<sup>ΔFlox</sup> | *Wapl*<sup>+/Flox</sup> |
| Observed | 0 | 20 |  | Observed | 7 | 6 |
| Expected | 10 | 10 |  | Expected | 6.5 | 6.5 |

### Looking for genetic interactions between *Wapl* and *Nipbl*

In Drosophila, a developmental defect caused by a *Wapl* dominant negative allele could be corrected by decreasing *Nipbl* gene dosage (*26*). This led us to hypothesize that decreasing *Wapl* levels in mice might correct developmental defects present in *Nipbl* mutants. To test this, we performed two independent crosses as described in Fig. 2, A and B. In Cross 1, we generated *Nipbl*^*-/+*^ animals in both *Wapl*^*+/+*^ and *Wapl*^*Δ/+*^ backgrounds (Fig. 2A). In Cross 2, we generated *Nipbl*^*-/+*^ mice in *Wapl*^*+/+*^, *Wapl*^*Flox/+*^, *Wapl*^*Δ/+*^, and in *Wapl*^*Δ/Flox*^ backgrounds (Fig. 2B). We assayed survival to weaning, embryonic growth, and brain transcriptomes to test if decreased *Wapl* gene function would ameliorate *Nipbl*^*-/+*^ phenotypes.

**Figure 2.**
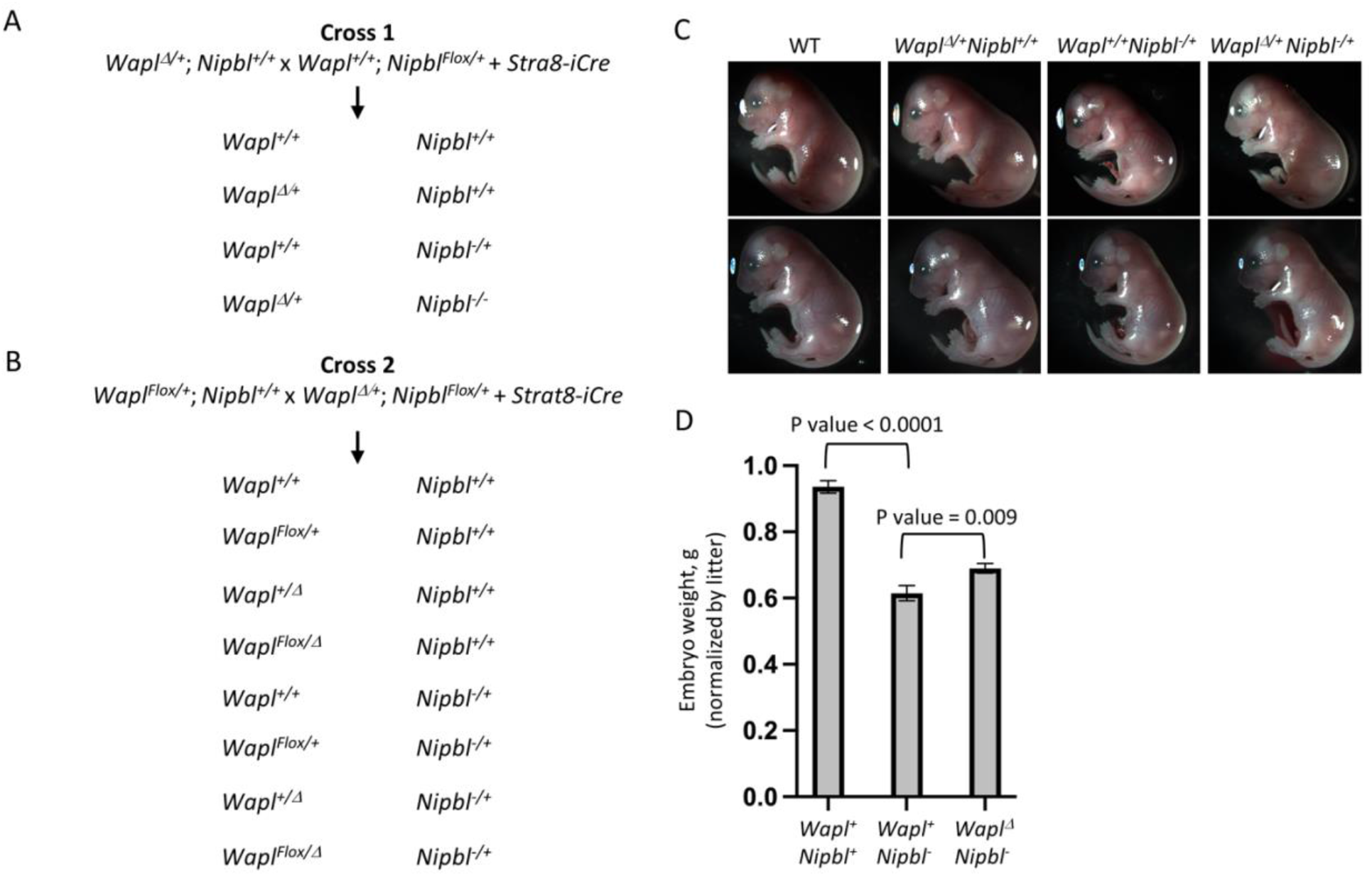
Generating Wapl^Δ/+^; Nipbl^-/+^ double heterozygotes along with wild type littermates and littermates that are heterozygous for loss of function mutations in either Wapl or Nipbl. ***A, B***. Crosses used to generate animals for this study. *Nipbl*^*-/+*^ mice die before weaning. Therefore, *Nipbl* mutants are maintained as *Nipbl*^*Flox/+*^ animals. The *Stra8-iCr*e transgene expresses Cre recombinase in pre-meiotic male germ cells. Thus *Nipbl*^*Flox/+*^ *Stra8-iCre* sires generate either *Nipbl*^*-*^ or *Nipbl*^*+*^ sperm. Because the *Stra8-iCre* transgene is expressed only in pre-meiotic germ cells and not in any other tissues, the maternally inherited *Wapl*^*Flox*^ allele in Cross 2 is not recombined in the progeny animals. **C**. Images of embryos from Cross 1 (top row) and Cross 2 (bottom row). **D**. *Nipbl* mutants show a strong growth deficiency that is partially compensated by reducing *Wapl* gene dosage. To account for the significant litter-to-litter variations in embryo size, we used a linear regression model that considered both genotype and litter as independent variables. In the analyses depicted here, *Wapl*^*Δ/+*^ and *Wapl*^*Flox/Δ*^ animals were pooled and we see 24% rescue of the growth defect (N = 56; P value = 0.009). When we excluded animals carrying a *Wapl*^*Flox*^ allele, we see a 19% rescue (N = 41, P value = 0.03).

### Reduced *Wapl* function does not rescue *Nipbl+/Nipbl-* postnatal lethality

We analyzed 79 progeny from Cross 1, genotyping weanlings at postnatal day 21 (Table 2). *Wapl* inheritance followed normal Mendelian patterns: 38 mice were *Wapl*^*Δ/+*^ and 41 mice were *Wapl*^*+/+*^ (c^2^ = 0.114, 1 degree of freedom; p = 0.74). However, no *Nipbl*^*-/+*^ animals were identified. We next analyzed 58 progeny from Cross 2 (Table 3). Again, *Wapl* inheritance followed the expected patterns, but no *Nipbl*^*-/+*^ animals were identified. Thus, neither the *Wapl*^*Δ/+*^ nor the *Wapl*^*Δ/Flox*^ backgrounds facilitated survival of *Nipbl*^*-/+*^ pups. Similarly, a *Nipbl*^*-/+*^ background did not permit survival of *Wapl*^*Δ/Flox*^ pups.

**Table 2.** Decreasing *Wapl* dosage does not rescue lethality in *Nipbl*^*-/+*^ mice. Genotypes of 79 weanlings were determined. Expected^1^, numbers of progeny expected assuming all genotypes survive; χ^2^ = 79.228 with 3 degrees of freedom; P value < 0.0001. Expected^2^, numbers of progeny expected assuming *Wapl* haploinsufficiency fully rescues lethality in *Nipbl*^*-/+*^ mice; χ^2^ = 39.671 with 2 degrees of freedom, P value < 0.0001. Expected^3^, numbers of progeny assuming *Wapl* haploinsufficiency cannot rescue lethality in *Nipbl*^*-/+*^ mice; c^2^ = 0.114 with 1 degree of freedom, P value = 0.7357.

| Cross: *Wapl*<sup>Δ/+</sup>; *Nipbl*<sup>+/+</sup> x *Wapl*<sup>+/+</sup>; *Nipbl*<sup>Flox/+</sup> *Stra8Cre* |  |  |  |  |
| --- | --- | --- | --- | --- |
| Genotype | *Wapl*<sup>Δ/+</sup> *Nipbl*<sup>+/+</sup> | *Wapl*<sup>+/+</sup> *Nipbl*<sup>+/+</sup> | *Wapl*<sup>Δ/+</sup> *Nipbl*<sup>-/+</sup> | *Wapl*<sup>+/+</sup> *Nipbl*<sup>-/+</sup> |
| Observed | 38 | 41 | 0 | 0 |
| Expected<sup>1</sup> | 19.75 | 19.75 | 19.75 | 19.75 |
| Expected<sup>2</sup> | 26.33 | 26.33 | 26.33 | 0 |
| Expected<sup>3</sup> | 39.5 | 39.5 | 0 | 0 |

**Table 3.** Decreasing *Wapl* gene dosage does not rescue lethality in *Nipbl*^*-/+*^ mice. Genotypes of 58 weanlings were determined. Expected^1^, numbers of progeny expected assuming all genotypes survive; χ^2^ = 79.228 with 3 degrees of freedom, P value < 0.0001. Expected^2^, numbers of progeny expected assuming *Wapl* haploinsufficiency fully rescues lethality in *Nipbl*^*-/+*^ mice; χ^2^ = 39.671 with 2 degrees of freedom, P value < 0.0001. Expected^3^, numbers of progeny assuming *Wapl* haploinsufficiency cannot rescue lethality in *Nipbl*^*-/+*^ mice; χ^2^ = 0.114 with 1 degree of freedom, P value = 0.7357.

| Cross: *Wapl*<sup>Flox/+</sup>; *Nipbl*<sup>+/+</sup> x *Wapl*<sup>Δ/+</sup>; *Nipbl*<sup>Flox/+</sup> *Stra8Cre* |  |  |  |  |  |  |  |  |
| --- | --- | --- | --- | --- | --- | --- | --- | --- |
| Genotypes | *Wapl*<sup>Flox/Δ</sup> *Nipbl*<sup>+/+</sup> | *Wapl*<sup>+/Δ</sup> *Nipbl*<sup>+/+</sup> | *Wapl*<sup>Flox/+</sup> *Nipbl*<sup>+/+</sup> | *Wapl*<sup>Flox/+</sup> *Nipbl*<sup>+/+</sup> | *Wapl*<sup>Flox/Δ</sup> *Nipbl*<sup>-/+</sup> | *Wapl*<sup>+/Δ</sup> *Nipbl*<sup>-/+</sup> | *Wapl*<sup>+/Δ</sup> *Nipbl*<sup>-/+</sup> | *Wapl*<sup>+/Δ</sup> *Nipbl*<sup>-/+</sup> |
| Observed | 0 | 19 | 20 | 19 | 0 | 0 | 0 | 0 |
| Expected<sup>1</sup> | 7.25 | 7.25 | 7.25 | 7.25 | 7.25 | 7.25 | 7.25 | 7.25 |
| Expected<sup>2</sup> | 0 | 11.6 | 11.6 | 11.6 | 11.6 | 11.6 | 0 | 0 |
| Expected<sup>3</sup> | 0 | 19.33 | 19.33 | 19.33 | 0 | 0 | 0 | 0 |

### Reduced *Wapl* function partially rescues *Nipbl*^*-/+*^ embryonic growth deficiency

Previous analyses showed that *Nipbl*^*-/+*^ embryos display a variety of developmental defects whose penetrance varies greatly from animal to animal (*14, 23*). One phenotype that is consistently observed in *Nipbl*^*-/+*^ embryos is significantly reduced growth. Therefore, we tested the effect of *Wapl*-deficiency on embryo weight in both *Nipbl*^*+/+*^ and *Nipbl*^*-/+*^ backgrounds.

We identified mice in proestrus or estrus and set up matings at 14:00. Mating pairs were separated by 07:30 the next day and embryos were collected between 11:00 and 13:00 on E17.5. With Cross 1, we generated 6 litters and 34 embryos. With Cross 2, we generated 9 litters and 62 embryos. See Fig. 2C for images of representative embryos. Embryo weights from these two crosses were collected and data from the two crosses were pooled for evaluation (Table S5).

To account for the significant litter-to-litter variations in embryo size (*28*) we analyzed the data by linear regression using a two-factor model that considered both genotype and litter as independent variables. As previously reported, *Nipbl*-deficiency leads to reduced embryo size (*Nipbl*^*+/+*^ = 0.936 g, *Nipbl*^*-/+*^ = 0.615 g or 34% decrease; P value = 2.95e-13) (Fig. 2D). In contrast, *Wapl*-deficiency has a modest effect (4.5% decrease) that is not statistically significant (P value = 0.14).

Our primary interest was in the effect of *Wapl* deficiency in a *Nipbl*^*-/+*^ background. When controlling for the effects of litter, double heterozygotes (*Wapl*^*Δ/+*^; *Nipbl*^*-/*+^ or *Wapl*^*Δ/Flox*^; *Nipbl*^*-/+*^) are significantly larger than *Nipbl*^*-/+*^ embryos (*Nipbl*^*-/+*^; *Wapl*^*Δ/+*^ = 0.690 g, *Nipbl*^*-/+*^ = 0.615 g or 12% increase; P value = 0.00932) (Fig. 2D). This means that 24% of the *Nipbl*^*-/+*^ growth phenotype was rescued by reducing *Wapl* gene dosage.

### *Wapl* RNA levels are insensitive to *Nipbl* heterozygosity and *Nipbl* RNA levels are insensitive to *Wapl* heterozygosity

We isolated total RNA from E17.5 embryonic brains and quantitated *Wapl* and *Nipbl* expression by qRT-PCR. Brains from mice heterozygous for the *Nipbl*^*-*^ allele show 50% loss of *Nipbl* RNA (P value < 0.001) (Fig. 3A). Similarly, brains from mice heterozygous for the *Wapl*^*Δ*^ allele show 50% reduction in *Wapl* RNA (P value < 0.001) (Fig. 3B). We saw no evidence that the *Wapl*^*Flox*^ allele alters *Wapl* RNA levels.

**Figure 3.**
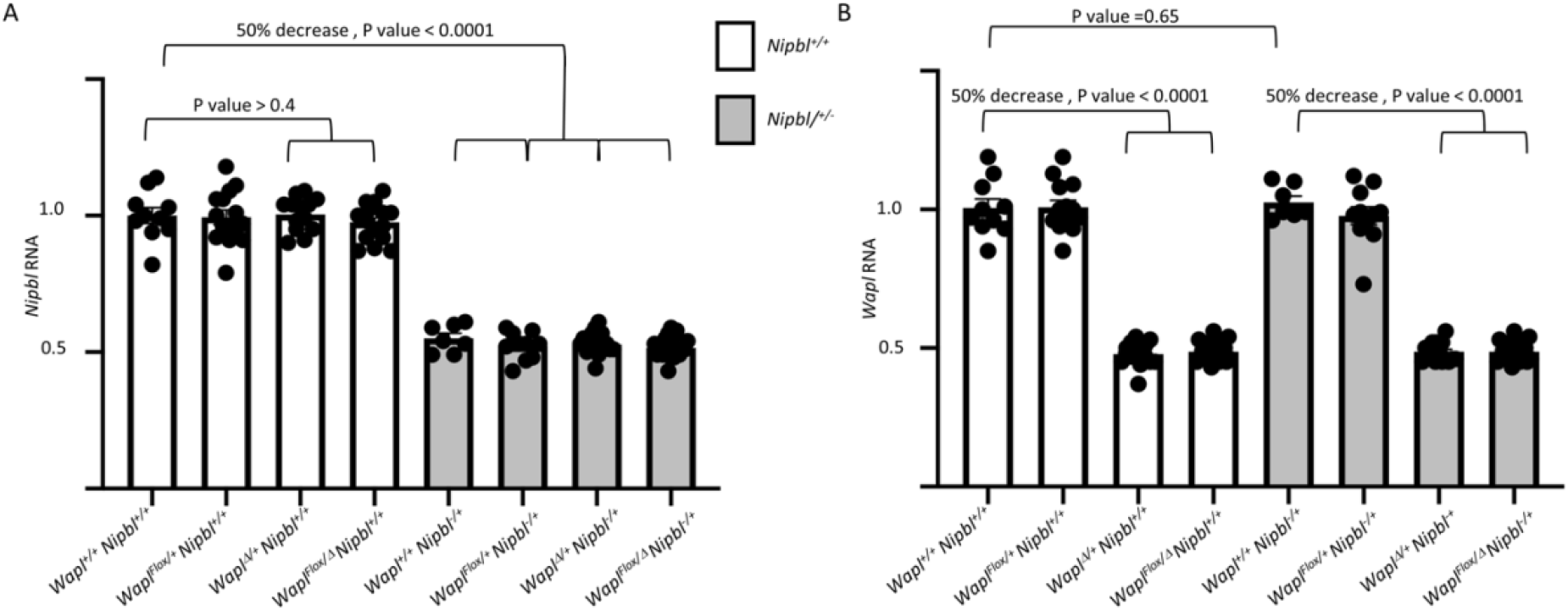
Wapl and Nipbl transcription. RNAs were isolated from E17.5 brains, analyzed by qRT-PCR, normalized to *GAPDH*, and then normalized to levels in wild type (*Wapl*^*+/+*^; *Nipbl*^*+/+*^) animals. **A**. *Wapl* RNA levels are reduced 2-fold in *Wapl*^*Δ/+*^ heterozygotes but are not affected by mutations in *Nipbl*. **B**. *Nipbl* RNA levels are reduced 2-fold in *Nipbl*^*+/-*^ heterozygotes but are not altered by mutations in *Wapl*.

Most importantly for this study, *Nipbl* levels are not altered by *Wapl* mutations and *Wapl* RNA levels are not altered by *Nipbl* mutations. This information was essential for designing and interpreting the transcriptome analyses described below.

### Transcriptome changes in *Nipbl*^*-/+*^ and *Wapl*^*Δ/+*^ embryonic brains

RNA-Seq data was generated from whole brain tissue derived from female E17.5 embryos. 22 samples were sequenced: 5 *Wapl*^*+/+*^; *Nipbl*^*+/+*^ (referred to as *WT*), 7 *Wapl*^*Δ/+*^, 4 *Nipbl*^-/+^, and 6 *Wapl*^Δ/+^; *Nipbl*^*-/+*^ double heterozygotes. After quality control assessment, one *Nipbl*^*-/+*^ sample was removed from further analysis due to poor read generation, while the remaining 21 samples were assessed to be of good quality (Fig. S1).

Principal component analysis (PCA) illustrates the segregation of replicates by genotype for *WT, Wapl*^*Δ/+*^, and *Nipbl*^*-/+*^ samples (Fig. 4A). However, double heterozygote replicates do not cluster. Note that two double heterozygote replicates co-localize with the *WT* samples. While *Nipbl*^*-/+*^ and *Wapl*^*Δ/+*^ replicates cleanly cluster away from *WT* samples, their replicates are more dispersed, indicating greater intragroup variability. This increased intragroup variability is consistent with the phenotypic variability observed in *Nipbl*^-/+^ mice, where the penetrance of many abnormal neurological phenotypes is less than 50% (*14*).

**Figure 4.**
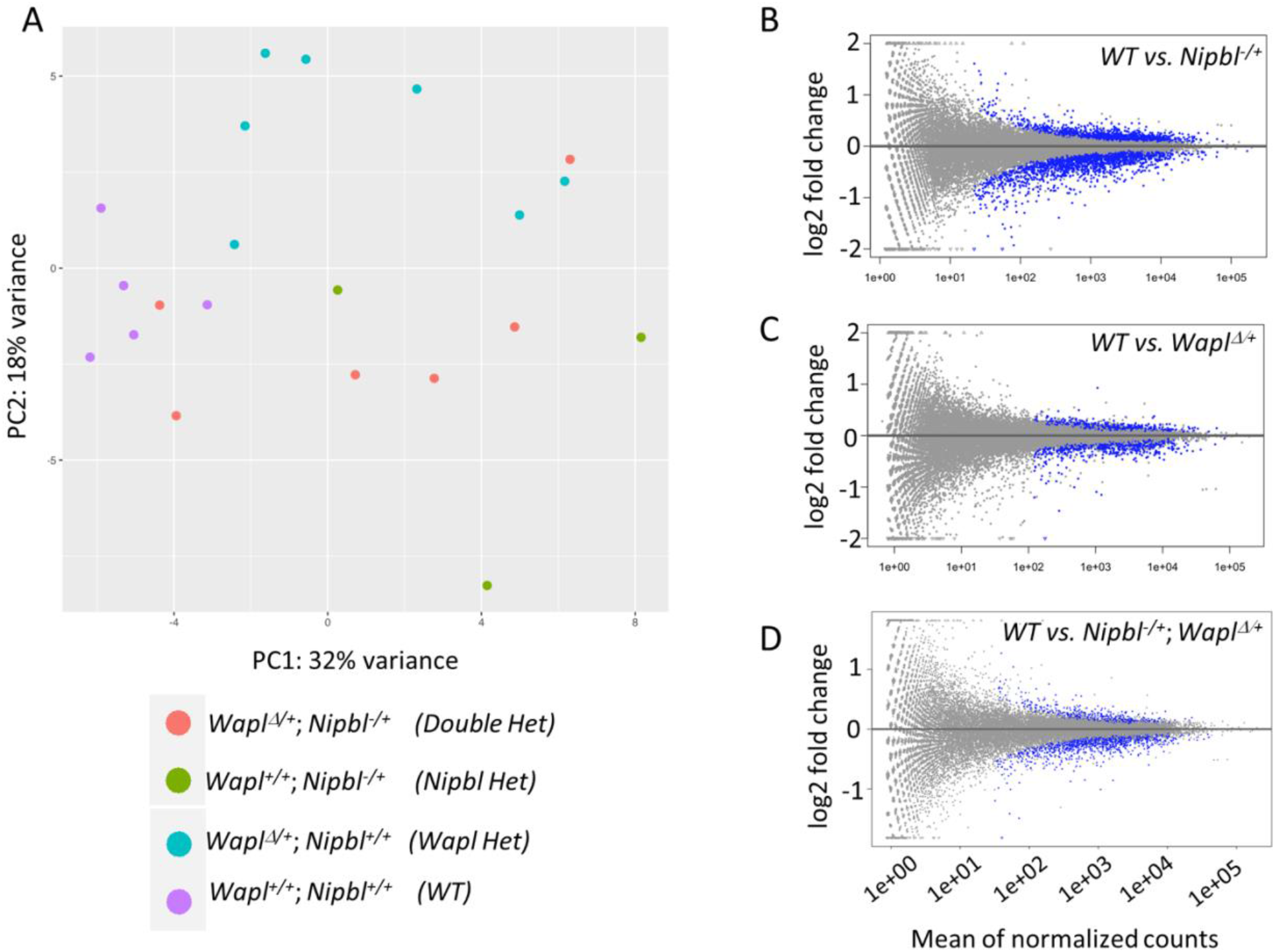
Transcriptome Analyses. **A**. PCA was performed as described in Methods. Wild type replicates (purple) are tightly clustered. *Nipbl*^*-/+*^ (green) and *Wapl*^*Δ/+*^ (blue) replicates show greater sample-to-sample variation but still cluster into distinct groups. In contrast, double heterozygote replicates (red) show high variability and individual samples cluster with both wild type and with individual heterozygotes. **B-D**. Differentially expressed genes. MA plots for *wild type* vs. *Nipbl*^*-/+*^ **(B)**, *wild type* vs. *Wapl*^*Δ/+*^ **(C)**, and *wild type vs. Nipbl*^*-/+*^; *Wapl*^*Δ/+*^ samples **(D)**. Blue dots denote genes that are differentially expressed. In **B, C**, and **D**, differentially expressed genes (DEGs) are normally distributed regarding their expression levels in wild type samples. Also, for **B, C**, and **D** the change in expression in mutant samples is typically modest: >98% of DEGs show fold changes of less than 2.

Differential expression analysis was performed by DESeq2 using an FDR-adjusted P value significance threshold of 0.10 (*29*). Comparing *WT* to *Nipbl*^*-/+*^ transcriptomes, 1535 genes were significantly upregulated, and 1971 genes were downregulated for a total of 3506 differentially expressed genes (DEGs) (Fig. 4B, Table S6). Of these DEGs, 3460 (98.7%) were dysregulated by less than 2-fold. Comparing *WT* to *Wapl*^*Δ/+*^ transcriptomes, a similar pattern of dysregulation was observed; 697 genes were significantly upregulated and 730 genes were significantly downregulated for a total of 1427 differentially expressed genes (Fig. 4C, Table S7). Of the 1427 DEGs, 1421 (99.6%) were dysregulated by less than 2-fold. Taken together, the transcriptomic dysregulation observed in both *Nipbl*^-/*+*^ and *Wapl*^*Δ/+*^ brains is consistent with the broad, low-effect transcriptomic dysregulation reported when studying mutations in cohesin related genes (*13, 30, 31*).

WAPL unloads and NIPBL loads cohesin onto chromosomes. If the transcriptional dysregulation observed in mutant heterozygotes stems from a disruption to cohesin function, one might expect a large set of shared DEGs that define loci where transcription is sensitive to cohesin structures. In fact, 851 genes are dysregulated in both *Wapl*^*Δ/+*^ and *Nipbl*^*-/+*^ mutants (P value < 0.0001, chi square of proportions) (Fig. 5A). Consistent with previous analyses in Drosophila (*32*), dysregulation in *Nipbl* and *Wapl* mutants is almost always in the same direction (99% of points are in the top-right and bottom-left quadrants of Fig. 5B).

**Figure 5.**
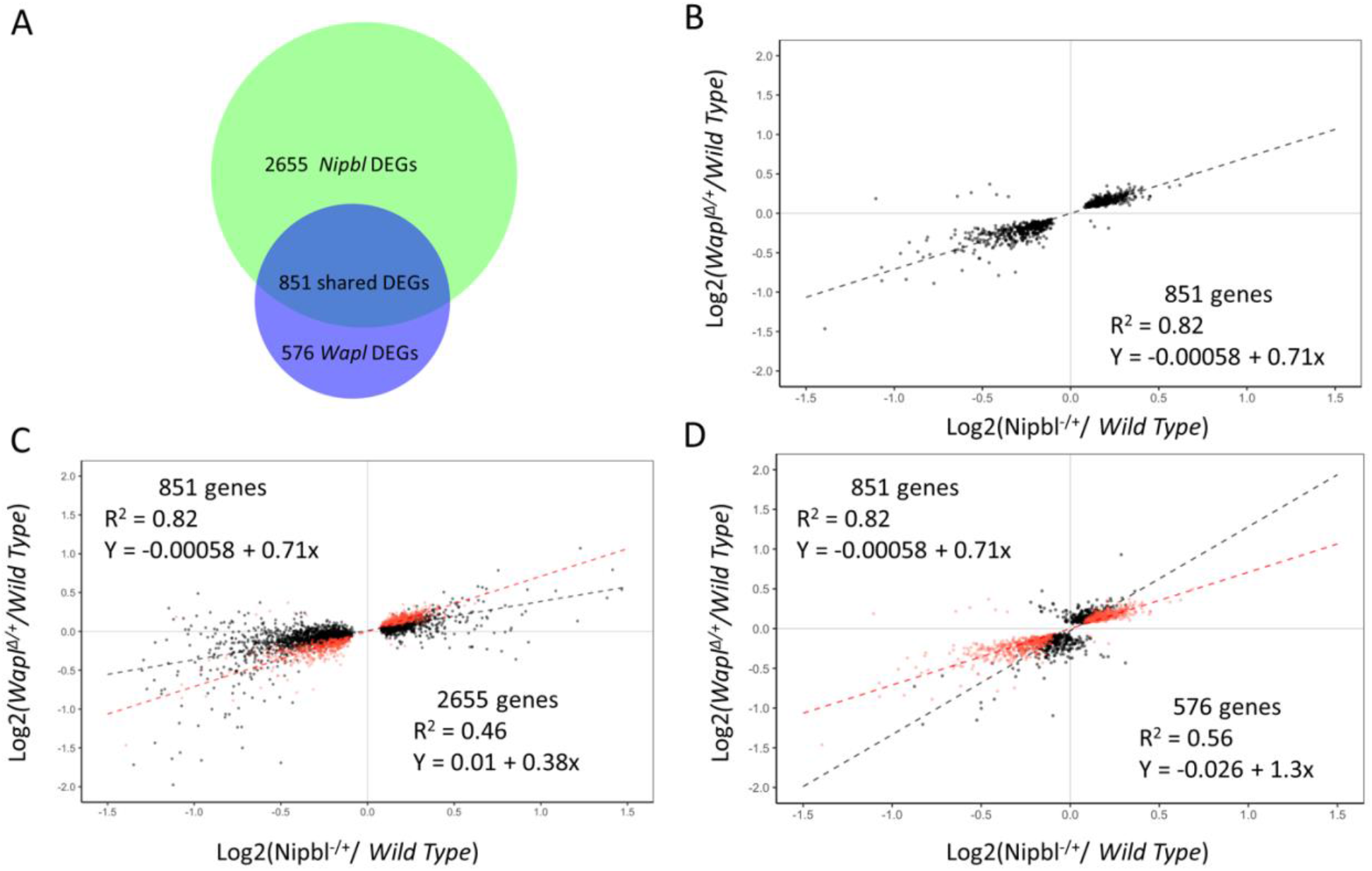
Overlapping transcriptome defects in Wapl^Δ/+^ and in Nipbl^-/+^ embryos. **A**. 3506 genes are dysregulated in *Nipbl*^*-/+*^ brains and 1427 genes are dysregulated in *Wapl*^*Δ/+*^ brains relative to wild type samples. 851 genes are dysregulated in both sample sets. The overlap is highly significant (P value < 0.0001, chi square of proportions). **B**. Linear regression analysis describing dysregulation phenotypes of 851 overlapping DEGs. Each gene (black dot) is plotted according to its dysregulation phenotype in *Nipbl*^*-/+*^ (x-axis) and *Wapl*^*Δ/+*^ (y-axis) samples. Note that dysregulation is almost always in the same direction and to a similar degree. Black dashes, Line of Best Fit; Coefficient of determination, R^2^ = 0.82. **C**. Scatterplot describing dysregulation phenotype in all 3506 *Nipbl* DEGs. Red dots show the 851 shared DEGs. Linear regression analyses were performed separately on 851 overlapping DEGs and also on the 2654 genes that were not identified as DEGs in *Wapl* mutants. **D**. Scatterplot describing the dysregulation phenotype in all 1427 *Wapl* DEGs. Red dots show the 851 shared DEGs. Linear regression analyses were performed separately on 851 overlapping DEGs and also on the 576 genes that were not identified as DEGs in *Nipbl* mutants. In the latter two figures, the red dashes represent the line of best fit for the 851 shared DEGs while the black dashes represent the 2655 & 576 non-shared DEGs.

The overlap in DEGs is large (25% of *Nipbl* DEGs and 60% of all *Wapl* DEGs) but we suspect it is still an underestimate of the overlap in transcriptional defects. Linear regression analyses suggest that most *Nipbl* DEGs show a trend toward dysregulation in *Wapl*^*Δ/+*^ samples, even if the *Wapl* effect was not statistically significant enough to score the gene as a *Wapl* DEG (89%, Fig. 5C). Similarly, most *Wapl* DEGs show a trend toward dysregulation in *Nipbl*^*-/+*^samples (88%, Fig. 5D).

### Decreased *Wapl* dosage partially rescues the transcriptome dysregulation observed in *Nipbl*^*-/+*^ mice

We focused next on analyses of the double mutant where there are only 1473 DEGs relative to wild type samples (Fig. 4C). If altered cohesin dynamics are responsible for the transcriptional changes observed in *Wapl*^*Δ*/*+*^ and *Nipbl*^*-/+*^ mutants, it is possible that correcting cohesin dynamics could restore the transcriptome towards *WT*. This idea led to the hypothesis that a *Wapl*^*Δ*/*+*^; *Nipbl*^*-/+*^ mutant may possess a more *WT*-like transcriptome than either a *Wapl*^*Δ*/*+*^ or a *Nipbl*^*-/+*^ mutant, for the double heterozygote would possess a *Wapl*:*Nipbl* dosage ratio like *WT*. To test whether the double heterozygote does, in fact, rescue the transcriptional dysregulation observed in *Nipbl*^*-/+*^ mutants, the 3506 genes differentially expressed between *WT* and *Nipbl*^*-/+*^ were clustered based on their expression levels in *WT, Nipbl*^*-/+*^, and *Wapl*^*Δ/+*^; *Nipbl*^*-/+*^ embryonic brains (Pantano, 2022). The unbiased clustering identified six classes of genes, which were annotated as complete rescue of upregulation (Group 1, 51 genes), partial rescue of upregulation (Group 2, 1404 genes), no rescue of upregulation (Group 3, 80 genes), complete rescue of downregulation (Group 4, 18 genes), partial rescue of downregulation (Group 5, 1825 genes), and no rescue of downregulation (128 genes) (Fig. 6A; Table S8). In total, four out of the six classes, containing 3298 of 3506 (94.1%) *Nipbl* DEGs were identified as exhibiting transcriptional rescue mediated by a reduction of *Wapl* dosage.

**Figure 6.**
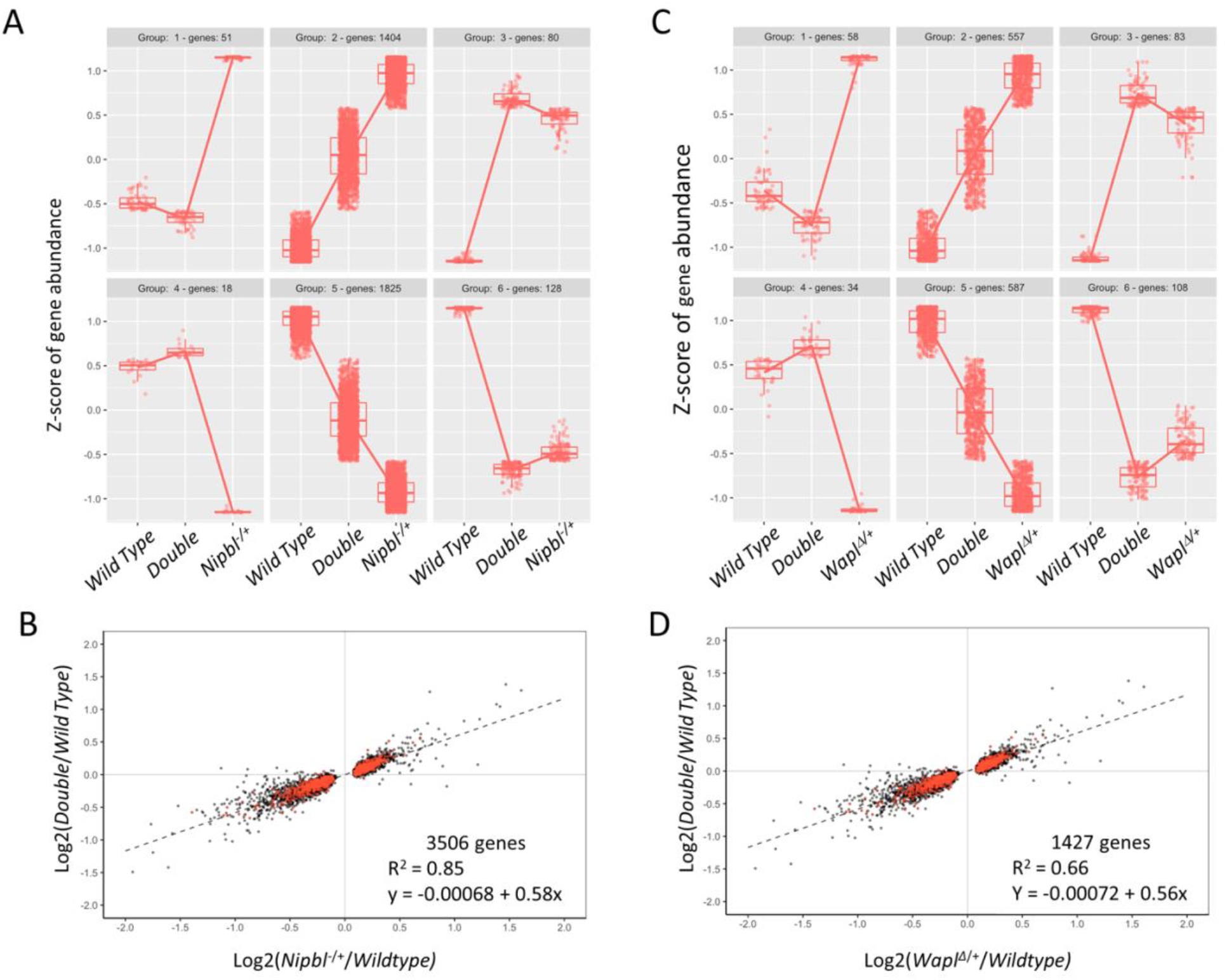
Rescue of transcriptome phenotypes in Wapl^Δ/+^; Nipbl^-/+^ double heterozygotes. **A, B**. Rescue of *Nipbl* transcriptome defects. **A**. 94% of the 3506 genes dysregulated in *Nipbl*^*-/+*^ brains are at least partially rescued by concomitant reduction in *Wapl* gene function. For each gene differentially expressed in *Nipbl* heterozygotes, normalized counts from wild type, double heterozygotes (*Wapl*^*Δ/+*^; *Nipbl*^*-/+*^), and *Nipbl*^*-/+*^ samples were analyzed using a DIANA clustering algorithm. **B**. Linear regression analyses. For each *Nipbl* DEG, the magnitude of dysregulation in *Nipbl* heterozygotes (x-axis) is plotted against the magnitude of dysregulation in double heterozygotes (y-axis). The regression line summarizes the overall effect of *Wapl* mutation. A slope of 1 would indicate no rescue while a slope of 0 would indicate complete rescue. Here the slope is 0.58 indicating a 42% rescue. The R^2^ of 0.85 demonstrates the consistency of rescue across the 3506 DEGs. Red dots represent genes that are also significantly dysregulated in *Wapl*^*Δ/+*^ heterozygotes. **C, D**. Rescue of *Wapl* transcriptome defects. **C**. 87% of the 1427 genes dysregulated in *Wapl*^*Δ/+*^ brains are at least partially rescued by concomitant reduction in *Nipbl* gene function. For each gene differentially expressed in *Wapl* heterozygotes, normalized counts from wild type, double heterozygotes (*Wapl*^*Δ/+*^; *Nipbl*^*-/*+^), and *Wapl*^*Δ/+*^ samples were analyzed using a DIANA clustering algorithm. **D**. Linear regression analyses. For each DEG, the magnitude of dysregulation in *Wapl* heterozygotes (x-axis) is plotted against the magnitude of dysregulation in double heterozygotes (y-axis). Here, the regression line summarizes the overall effect of *Nipbl* mutation and a slope of 0.56 denotes a 44% rescue. Red dots represent genes that are also significantly dysregulated in *Nipbl*^*-/+*^ heterozygotes.

Effect ratios were compared between *Nipbl*^*-/+*^ and *Wapl*^*Δ/+*^; *Nipbl*^*-/+*^ mutants to further characterize the transcriptional rescue phenotype (Fig. 6B; Table S9). Plotting the *Nipbl*^*-/+*^ / *WT* effect ratio on the x-axis and the *Wapl*^*Δ/+*^; *Nipbl*^*-/+*^ / *WT* effect ratio on the y-axis for each of the 3506 dysregulated genes enabled linear regression to be performed to estimate both the average magnitude of transcriptional rescue and the consistency of transcriptional rescue across the entire dysregulated gene set. Dysregulation in *Wapl*^*Δ/+*^; *Nipbl*^-/+^ mutants was 58% as severe as the dysregulation observed in *Nipbl*^*-/+*^ mutants. (See the slope of the regression line in Fig. 6B.) Importantly, this reduction in dysregulation was consistent across the gene set, as demonstrated by a coefficient of determination value (R^2^) of 0.85. These results demonstrate the ability of decreased *Wapl* dosage to rescue dysregulation caused by reductions in *Nipbl* dosage.

Similarly, the dysregulation observed in the *Wapl*^*Δ/+*^; *Nipbl*^*-/+*^ mutants was less severe than the dysregulation observed in the *Wapl*^*Δ/+*^ mutants. Of the 1427 dysregulated genes identified in the *Wapl*^*Δ/+*^ embryo, 1236 (86.6%) were clustered into 4 groups representing transcriptional rescue phenotypes: complete rescue of upregulation, (Group 1, 58 genes), partial rescue of upregulation (Group 2, 557 genes), complete rescue of downregulation (Group 4, 34 genes), and partial rescue of downregulation (Group 5, 587 genes) (Fig. 6C; Table S10).

Executing linear regression on plotted effect ratios reveled that dysregulation in *Wapl*^*Δ/+*^; *Nipbl*^*-/+*^ mutants was only 56% as severe as dysregulation in *Wapl*^*Δ/+*^ mutants on average (Fig. 6D; Table S9). A coefficient of determination value (R^2^) of 0.66 indicates a modest consistency in the magnitude of transcriptional rescue observed.

In Fig. 7 we analyze rescue of the 851 shared DEGs. These results emphasize that transcriptional rescue in the double heterozygotes occurs even when dysregulation in *Nipbl*^*-/+*^ and in *Wapl*^*Δ/+*^ samples are in the same direction. 40% of shared DEGs are upregulated in both *Nipbl* and *Wapl* heterozygotes and then partially rescued in the double mutant (Group 1). Similarly, 39% of shared DEGs are downregulated in both heterozygotes and then partially rescued in the double mutant (Group 4) (Fig. 7A, Table S11). On average we see a 41% rescue of *Nipbl* dysregulation (R^2^ = 0.91) (Fig. 7B) and a 32% rescue of *Wapl* dysregulation (R^2^ = 0.74) (Fig. 7C).

**Figure 7.**
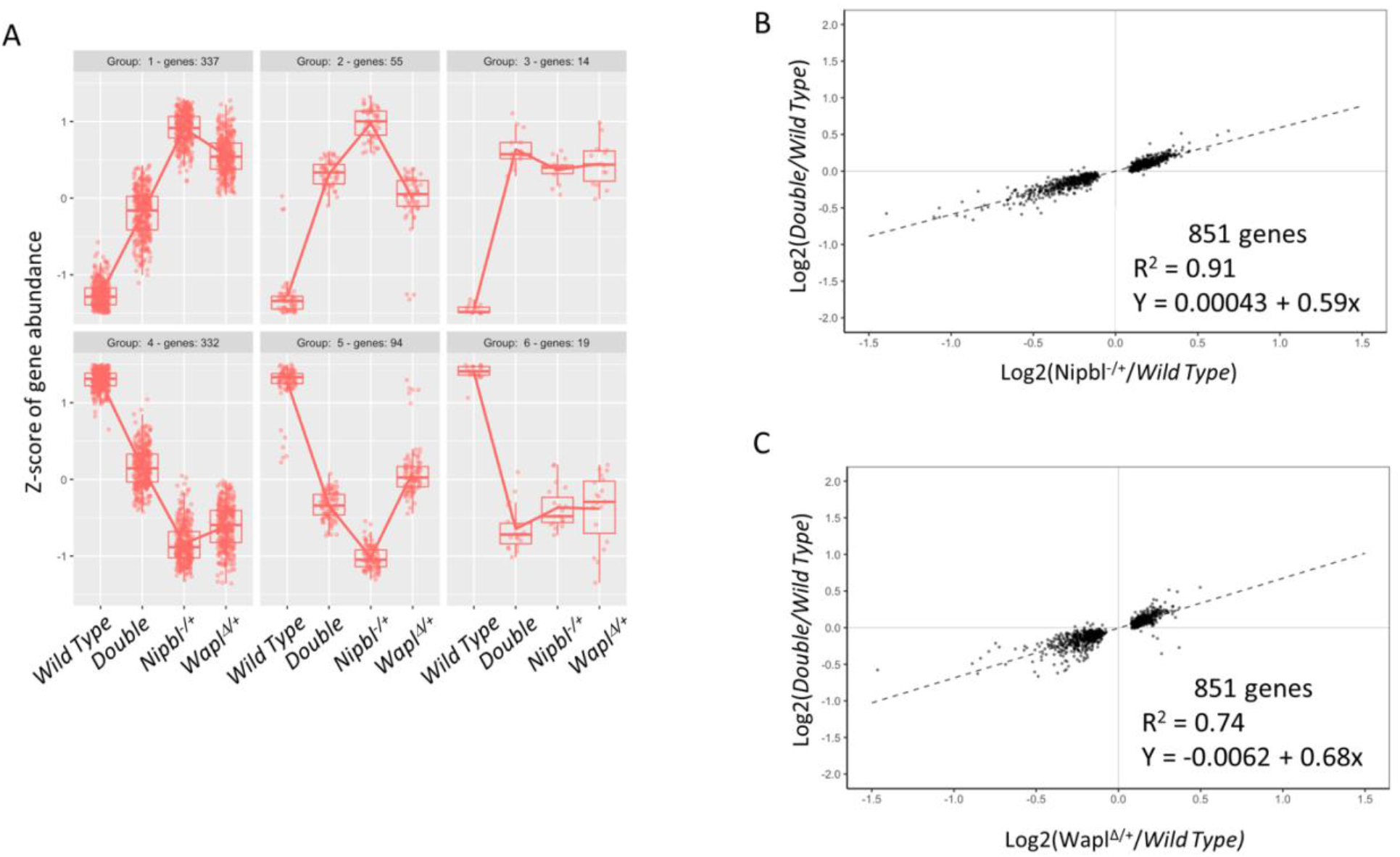
Rescue of overlapping transcriptome defects in Wapl^Δ/+^; Nipbl^-/+^ double heterozygotes. **A**. 79% of the 851 genes misexpressed in both *Nipbl*^*-/+*^ and *Wapl*^*Δ/+*^ brains are at least partially rescued by concomitant reduction in either *Nipbl* or *Wapl* gene function (Group 1 & 4). For each gene differentially expressed in both *Nipbl* and *Wapl* heterozygotes, normalized counts from wild type, double heterozygotes (*Wapl*^*Δ/+*^; *Nipbl*^*-/*+^), *Nipbl*^*-/+*^, and *Wapl*^*Δ/+*^ samples were analyzed using a DIANA clustering algorithm. **B**. Linear regression analyses. For each overlapping DEG, the magnitude of dysregulation in *Nipbl* heterozygotes (x-axis) is plotted against the magnitude of dysregulation in double heterozygotes (y-axis). The regression line summarizes the overall effect of *Wapl* mutation. A slope of 1 would indicate no rescue while a slope of 0 would indicate complete rescue. Here the slope is 0.59 indicating a 41% rescue. The R^2^ of 0.91 demonstrates the consistency of rescue across the 851 DEGs. **C**. For each overlapping DEG, the magnitude of dysregulation in *Wapl* heterozygotes (x-axis) is plotted against the magnitude of dysregulation in double heterozygotes (y-axis). The regression line summarizes the overall effect of *Nipbl* mutation. Here the slope is 0.68 indicating a 32% rescue. The R^2^ of 0.74 demonstrates the consistency of rescue across the 851 DEGs.

## Discussion

In this mouse study we generated a novel conditional allele of *Wapl* and analyzed the effects of reducing *Wapl* gene dosage in *Nipbl*^*+/+*^ and in *Nipbl*^*-/+*^ backgrounds. Mouse development is sensitive to *Wapl* gene dosage. *Wapl*^*Δ/+*^ mice are viable and fertile, but *Wapl*^*Δ/Δ*^ embryos die even before implantation. *Wapl*^*Δ/Flox*^ (like *Nipbl*^*-/+*^) embryos are viable at E17.5 but die soon after birth and are absent at weaning.

*Wapl*^*Δ/+*^ brains display dysregulation of >1400 genes but the effect at a large majority of loci is less than 2-fold. This pattern is typical for cohesinopathy patients and for cohesinopathy models in mouse, zebrafish, and Drosophila, and is the same pattern we saw in our study in *Nipbl*^*-/+*^ brains. Dysregulation in *Wapl*^*Δ*^ and in *Nipbl*^*-*^ heterozygotes was not only similar in overall pattern but also in the identity of affected genes. Among *Wapl* DEGs, 60% are also identified as *Nipbl* DEGs and are almost always dysregulated in the same direction and to a similar degree. The fact that both *Nipbl* and *Wapl* depletion leads to such similar defects is consistent with models that stress the dynamic nature of cohesin localization on the chromosome (*4, 32*).

Most interestingly, transcriptome defects are significantly rescued in double heterozygotes. 94% of *Nipbl* DEGs and 87% of *Wapl* DEGs show at least partial rescue in the *Wapl*^*Δ/+*^; *Nipbl*^*-/+*^ brains, and there are no clear examples where dysregulation is exacerbated by the double mutation. This is especially intriguing since dysregulation in the single mutants is almost always in the same direction. Thus, *Wapl*/*Nipbl* interactions are a paradigm where two wrongs do make a right. Our data are in agreement with other recent findings that co-depletion of WAPL and NIBPL rescued cell proliferation and transcriptional defects in mammalian cells (*4*) and where reduced *Nipbl* gene function rescued developmental defects associated with a dominant-negative *Wapl* allele in Drosophila (*26*). Altogether these results indicate that the correct balance of cohesin loading and unloading activities, rather than the absolute amounts of WAPL and NIPBL, are most critical.

## Materials and Methods

### Mice

We generated mice carrying the *Wapl*^*Flox*^ allele in a two-step process. In step 1, mouse embryonic stem cells (R1 line, 129SV) were electroporated with linearized plasmid pMVW1. pMVW1 includes a 1.8 kb 5’ homology flank and a 4.1 kb 3’ homology flank to direct insertion of a 1.9 kb *NeoR* cassette (flanked with *Frt* elements) plus a 4.2 kb BamHI-XbaI fragment that carries *Wapl* sequences -517 bp to +3641 bp (relative to the main transcriptional start site) that are flanked with *loxP* sequences inserted as direct repeats. The linearized plasmid also included a 2.8 kb *Diphtheria Toxin-A* gene for negative selection. G418 resistant colonies were isolated and screened by using one primer from outside the 5’ flank (5’- ACCCGGTAGAATTGACCTGC) and one primer internal to the *NeoR* cassette (5’- GAGGAGGACAGTCTAGGGCA) to identify a 2067 bp band. Homologous recombination on the 3’ end was confirmed by amplification using one primer from outside the 3’ flank (5’- GATGTTCCTATAAGCCAAGAAGGC) and one primer from inside the 3’ *loxP* site (5’- GCAAAACAACCCCTCACTCC) that was followed by nested PCR (5’- GATGTTCCTATAAGCCAAGAAGGC and 5’-AGGGTGCTAATGAGATGGCTC) to identify a 236 bp band. Targeted clones were injected into C57BL/6J blastocysts and chimeric founders were bred to C57BL/6J females to establish the *Wapl*^*Flox*^*+NeoR* line. In step 2, *Wapl*^*Flox*^*+NeoR* heterozygotes were crossed to *ROSA26:FLPe* knock in transgenic females (JAX #003946) to remove the *NeoR* cassette via Flp-recombinase-mediated site-specific recombination and thus generate the *Wapl*^*Flox*^ mouse (Fig. 1). Animals were backcrossed to C57BL/6J at least 4 times before use in this study.

We generated the *Wapl*^*Δ*^ allele by crossing *Wapl*^*Flox*^ heterozygotes with *E2a-Cre* transgenic females (JAX #003724) to remove the 4.1 bp fragment that includes the *Wapl* promoter, exon 1, intron 1, exon 2, and the first 94 bp of intron 2. Animals were backcrossed to C56BL/6J at least 4 more times before use in this study.

Genotypes were determined by PCR analysis of gDNAs extracted from ear punch samples. For *Wapl*^*Flox*^ genotyping we used a two-primer assay (5’- GATGTTCCTATAAGCCAAGAAGGC and 5’- AGGGTGCTAATGAGATGGCTC) that yields bands of 208 and 242 bp that represent *Wapl*^*+*^ and *Wapl*^*Flox*^ alleles, respectively. For *Wapl*^*Δ*^ genotyping, we used a three-primer assay (5’-AGGTAGGGGACAGAACTCCG, 5’- AGAGAGCCAACGCAGGTAAA, and 5’- AACGCAAGCCTAGCAACCT) that yields bands of 145 and 335 bp that represent *Wapl*^*+*^ and *Wapl*^*Δ*^ alleles, respectively. This three-primer assay can also detect the *Wapl*^*Flox*^ allele (291 bp). For *Nipbl* genotyping, we followed the protocol outlined in Santos et al. using primers B045, B048, and B050 (*23*). For *Strat8-iCre* genotyping (*33*), we followed the protocol provided by Jackson Labs (JAX #017490).

For timed matings, we identified mice in proestrus or estrus by cytological evaluation of vaginal lavage samples (*34*). Mating pairs were set up in the early afternoon and then separated early the next morning.

All mouse studies were performed according to NIH and PHS guidelines and only after protocols were approved by the *Eunice Kennedy Shriver* National Institutes of Child Health and Human Development Animal Care and Use Committee.

### RNA Analyses by qRT-PCR

RNAs were isolated from snap-frozen tissue samples using Tri-Pure (Sigma-Aldrich 11667165001) and RNeasy Micro Kit (Qiagen 74004), analyzed using a Thermo Fisher NANODROP 2000c to evaluate purity and yield, and then stored at -70°C.

For qRT-PCR, cDNA samples were prepared with and without reverse transcriptase using random hexamer primers (Roche 04 887 352 001). cDNAs were analyzed using SYBR Green (Roche 04 887 352 001) on the Roche Light Cycler 480 II (45 cycles with annealing at 60°C) using primers for *Wapl* (5’-AGAGAGTGTAACAGTGCATAATCC, 5’- ACTGCTGAATCAGGTCTTCACA), *Nipbl* (5’-CTGATGTGGTTGCAGCATGT, 5’- TGAGTACAAGCTTTCTTCACAGGT), and *Gapdh* (5’-TCAATGAAGGGGTCGTTGAT, 5’- CGTCCCGTAGACAAAATGGT. Assay specificity was demonstrated by melting curve analyses and gel electrophoresis. Statistical significance was evaluated using the student’s T test.

### RNA-Seq Library preparation and sequencing

Total RNA was extracted from snap-frozen, whole brain tissue of female, E17.5 embryos using Tri-Pure isolation reagent (Sigma-Aldrich 11667165001) and Qiagen RNeasy Micro kit (Qiagen 74004) with on-column DNase I treatment (Qiagen 74004). Thermo Fisher NANODROP 2000c was used to evaluate purity and yield, and RNAs were stored at -70°C. Samples with RNA integrity numbers > 9.0 as determined using an Agilent BioAnalyzer were purified by oligo-d(T). Libraries were prepared using RNA Sample Prep V2 kit (Illumina) and sequenced on an Illumina HiSeq2500 platform. 101 bp paired-end reads were trimmed to remove adapters using cutAdapt v2.4 and mapped to the mouse genome mm10 (GRCm38.p6) using STAR v2.5.3a (*35*). The mapped reads were then counted using the featureCounts command of subread v1.6.4 (*36*). Sequencing and alignment quality was assessed using MultiQC v1.9 (*37*).

### Differential expression analysis

Count normalization and differential expression analyses were performed using the DESeq2 package in R (*29*). Specifically, WT gene expression was compared pairwise to gene expression of each of the mutant genotypes. Prior to differential expression testing, genes with fewer than 10 normalized counts summed across all samples were removed. After differential expression testing, genes with an FDR adjusted P value < 0.10 were called as differentially expressed and subjected to further analyses. The principal component analysis was performed using DESeq2’s plotPCA function using normalized, variance stabilizing transformed counts. The BioVenn R package was utilized to visualize the overlap between DEG lists (*38*).

### Identification of transcriptional rescue

To identify genotype-dependent patterns of gene expression, significant genes were clustered and visualized using the degPatterns function from the DEGreport R package (Pantano, 2022). For each gene, degPatterns first averaged the normalized counts across replicates within each genotype group and then implemented a DIANA clustering algorithm to group genes into clusters based on similar shifts in gene expression across genotypes. For visualization, expression values were Z-score transformed.

To quantify the magnitude and consistency of transcriptional rescue for all genes called as differentially expressed, log2 transformed effect ratios were calculated where *WT* gene expression was the common denominator to 1) *Nipbl*^-/+^ gene expression, 2) *Wapl*^*Δ/+*^ gene expression, and 3) *Wapl*^*Δ/+*^; *Nipbl*^*-/+*^ gene expression. Here, gene expression is equivalent to the normalized counts of a given gene averaged across the replicates of a given genotype. Effect ratios for differentially expressed genes were then plotted, comparing the magnitude of dysregulation imposed by the single heterozygotes – log2(*Nipbl*^*-/+*^ / *WT*) or log2(*Wapl*^*Δ/+*^ / *WT*) – to the magnitude of dysregulation imposed by the double heterozygotes – log2(*Wapl*^*Δ/+*^; *Nipbl*^*-/+*^ / *WT*). Simple linear regression was then performed to obtain a regression line, the slope of which was used to summarize the difference in dysregulation imposed by the plotted genotypes. Note that by plotting log2(*Wapl*^*Δ/+*^; *Nipbl*^-/+^ / *WT*) on the y-axis and either log2(*Nipbl*^*-/+*^ / *WT*) or log2(*Wapl*^*Δ/+*^ / *WT*) on the x-axis, slopes < 1 signify a lessened dysregulation of gene expression in the double heterozygotes. Additionally, coefficient of determination (R^2^) values were obtained from the simple linear regression, providing an estimation of the consistency of the observed pattern. Because the plots in Fig. 6 show effects in the double mutants on genes that had been selected based on their being significantly different from wild type in one or both single mutants, it seemed possible that some of the observed rescue was partly due to statistical regression to the mean or by false positives in the DEG lists. To control for this possibility, we repeated these analyses adding the 1473 genes differentially expressed in double mutants and saw no significant change in the slopes of the regression lines obtained from those shown in Fig. 6.

## Acknowledgments

We thank Alex Grinberg for generating the *Wapl*^*Flox*^ mouse line. We thank Jeanne Yimdjo and Victoria Biggs for animal husbandry. We thank the NICHD Molecular Genetics Core for library construction and sequencing used to generate the RNA-seq datasets.

## Author contributions

Conceptualization: CJT, JAK, KP

Methodology: CMK, CJT, AM, MTVW, ALC, JAK, KP

Software: CMK, AM

Investigation: CMK, CJT, MTVW, CMG, JN, KP

Formal Analysis: CMK, CJT, AM, CMG, ADL, JAK, KP

Visualization: CMK, CJT, AM, KP

Writing-original draft: CMK, JAK, KP

Writing-review and editing: AM, ALC, ADL

## Data and materials availability

RNA-seq data have been deposited in the National Center for Biotechnology Information Gene Expression Omnibus database under accession code GSE203014. All other data needed to evaluate the conclusions in the manuscript are present in the paper and/or the Supplementary Materials. The novel *Wapl* mouse lines can be obtained from KP.

