## Supplemental Figure 1 for "Decreasing *Wapl* dosage partially corrects transcriptome phenotypes in *Nipbl*-/+ embryonic mouse brain"

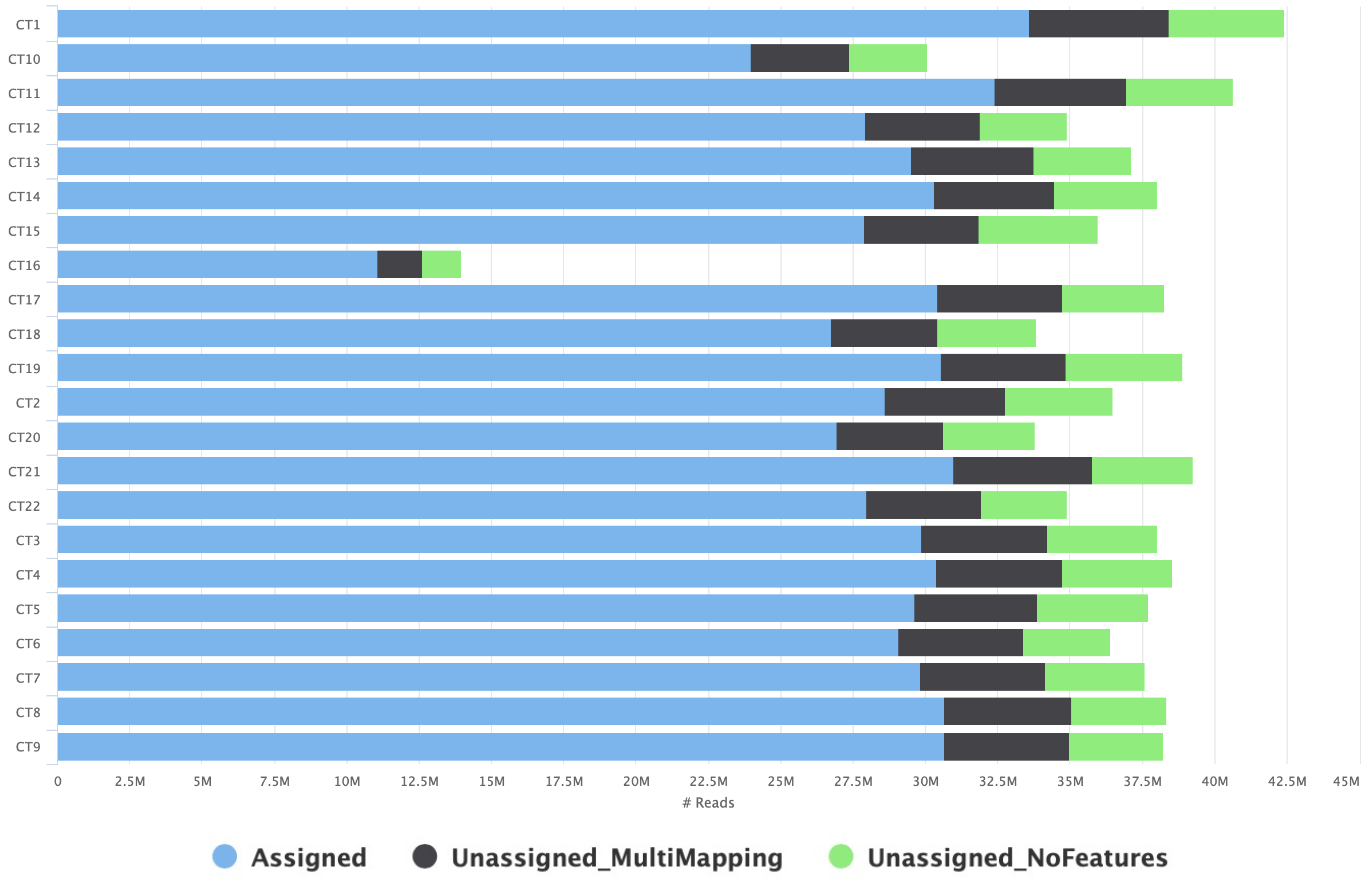


***Figure S1. RNA sequencing quality control assessment.*** RNAs were isolated from E17.5 brains and poly(A) molecules were sequenced as described in Methods. Sequencing quality was analyzed using MultiQC software. This graph depicts the number of uniquely mapped reads for each sample. Sample CT16, from a *Nipbl*$^{-/+}$ animal, was excluded from further analyses based on the failure to generate sufficient reads.
